# Intestinal microbiota modulates adrenomedullary response through Nod1 sensing in chromaffin cells

**DOI:** 10.1101/2020.10.02.323543

**Authors:** Chen Xiang, Peihua Chen, Qin Zhang, Yinghui Li, Ying Pan, Wenchun Xie, Jianyuan Sun, Zhihua Liu

## Abstract

The intestinal microbiota closely interacts with the neuroendocrine system and exerts profound effects on host physiology. Many of the molecular mechanisms underlying these interactions await discovery. Here we report that Nod1 ligand derived from intestinal bacteria directly modulates catecholamine storage and secretion in mouse adrenal chromaffin cells. The cytosolic peptidoglycan receptor Nod1 is involved in chromogranin A (CHGA) retention in dense core granules (DCGs) in chromaffin cells in a cell-autonomous manner. Mechanistically, upon recognizing its cognate ligands, Nod1 localizes to DCGs, and recruits Rab2a, which is critical for CHGA retention in DCGs. Loss of Nod1 ligands leads to a profound defect in epinephrine storage in chromaffin cells, and subsequently, less secretion upon stimulation. The intestine-adrenal medulla crosstalk bridged by Nod1 ligands modulates adrenal medullary responses during the immobilization-induced stress response in mice. Thus, our study uncovers a mechanism by which intestinal microbes directly modulate a major pathway in response to stress, which may provide further understanding of the gut-brain axis.

## Introduction

Accumulating evidence has shown that microbes residing in the intestine are deeply intertwined with diverse aspects of host physiology, ranging from metabolism, immune system development and immune responses, to neuronal development and activity (*Blander, Longman, Iliev, Sonnenberg, & Artis, 2017; Fung, Olson, & Hsiao, 2017; Nicholson et al., 2012; Rooks & Garrett, 2016*). It is emerging that the neuroendocrine system closely interacts with the gut microbiota (*Farzi, Frohlich, & Holzer, 2018*). Gut dysbiosis leads to profound alterations in neuroendocrine, neurochemical and behavioral parameters (*Crumeyrolle-Arias et al., 2014; Foster & McVey Neufeld, 2013; Rogers et al., 2016; Zheng et al., 2016*). Conversely, neuroendocrine can play a significant role in the gastrointestinal physiology, and malfunctioning neuroendocrine contributes to the pathogenesis of gastrointestinal disorders, including inflammatory bowel diseases (*Brinkman, Ten Hove, Vervoordeldonk, Luyer, & de Jonge, 2019; El-Salhy, Solomon, Hausken, Gilja, & Hatlebakk, 2017*). Uncovering the molecular mechanisms underlying such crosstalk between microbes and the neuroendocrine system is of importance to understand the role of the microbiota in host health and disease.

The neuroendocrine system controls vital processes in response to acute or chronic stress, which has been critical for host survival during the long history of evolution. The sympatho-adrenal-medullary (SAM) and hypothalamic-pituitary-adrenocortical (HPA) axes are the primary systems involved in the stress response. The SAM axis was first described by Cannon and colleagues to be involved in acute stress response, generally known as the “fight-or-flight” response (*Cannon, 1914*). The acute stress response involves increased circulating levels of adrenaline (primarily from the adrenal medulla) and noradrenaline (primarily from sympathetic nerves), increased heart rate and force of contraction, peripheral vasoconstriction, and energy mobilization (*Tank & Lee Wong, 2015*). For the HPA axis, stressors lead to secretion of corticotropin-releasing hormone (CRH) from the paraventricular nucleus (PVN) of the hypothalamus, and CRH acts on the pituitary gland, resulting in release of adrenocorticotrophic hormone (ACTH), which in turn cause the adrenal cortex to release the stress hormone cortisol (*Tsigos & Chrousos, 2002*). The HPA axis and the sympathetic system act in a complementary manner throughout the body during stress. Cortisol, together with epinephrine and norepinephrine, prepares the body for the stress response, which is characterized by gluconeogenesis, increased blood sugar, body temperature change, decreased immune function, and increased metabolism of fat and protein (*Khani & Tayek, 2001; Padgett & Glaser, 2003; Ste Marie & Palmiter, 2003; Tank & Lee Wong, 2015*).

It is interesting to know whether the intestinal microbiota may impact stress responses. Indeed, the effect of microbiota on HPA axis activation has been extensively examined. Absence of microbiota (germ-free status) in rodents leads to exaggerated activation of the HPA axis associated with elevated ACTH and corticosterone levels in response to stress, accompanied with anti-anxiety and anti-depression behavior changes (*Diaz Heijtz et al., 2011; Huo et al., 2017; Sudo et al., 2004; Zheng et al., 2016*). Of note, it is also reported that such behavior changes may depend on genetic background and husbandry (*Farzi et al., 2018*). Mechanistically, it has been shown that the microbiota regulates the expression of critical HPA axis genes both directly and indirectly (*Farzi et al., 2015; Luo et al., 2018; Shanks, Larocque, & Meaney, 1995*). One of the mechanisms may involve innate immune receptors, such as Toll-like receptors (TLRs). For instance, acute TLR4 activation triggers the HPA axis, likely through the action of cytokines (*Liu, Buisman-Pijlman, & Hutchinson, 2014; Zacharowski et al., 2006*). TLR2-deficient mice also exhibit impaired adrenal corticosterone release after inflammatory stress induced by bacterial cell wall compounds (*Bornstein et al., 2004*). In comparison, the effect of intestinal microbiota on the SAM pathway has not been extensively investigated. One previous finding found that the absence of gut microbial colonization selectively impaired catecholamine response to hypoglycemic stress in mice.(*Giri, Hu, La Gamma, & Nankova, 2019*)

In the SAM pathway, upon stimulation from sympathetic nerves, adrenal chromaffin cells located in the adrenal medulla secrete catecholamines (mainly epinephrine), which are the hormones necessary in the fight-or-flight response (*Eisenhofer, Kopin, & Goldstein, 2004*). Adrenal chromaffin cells convert tyrosine to catecholamines in a complex multi-step process (*Berends et al., 2019*). The catecholamines are then stored and bound to chromogranin A (CHGA) in dense core granules (DCGs) (*Jirout et al., 2010; Mahapatra et al., 2004; Videen, Mezger, Chang, & O’Connor, 1992*). CHGA is involved in DCG biogenesis by preventing degradation of granule proteins and packing synthetic catecholamines in DCGs (*Kim, Zhang, Sun, Wu, & Loh, 2005*). CHGA deficiency led to a 30-40% reduction in catecholamine secretion from adrenal chromaffin cells in mice (*Pasqua et al., 2016*). Similarly, knockdown of CHGA in PC-12 cells, a neuroendocrine cell line, decreased the number of DCGs and impaired DCG formation (*Kim, Tao-Cheng, Eiden, & Loh, 2001*). Thus, CHGA plays a key role in epinephrine storage in chromaffin cells.

Nucleotide-binding oligomerization domain 1 (Nod1), together with Nod2, as classic pattern recognition receptors (PRRs), play a key role in innate immunity responses by recognizing conserved microbial molecules from invading pathogens (*Chamaillard et al., 2003; Mogensen, 2009*). Studies have found that Nod1 and Nod2 are involved in mediating the gut-brain crosstalk. Mice deficient in both Nod1 and Nod2 (NodDKO) exhibited elevated anxiety levels in the context of a hyperactive HPA axis (*Schneider, Zhang, Barrett, & Gareau, 2016*). Specific deletion of Nod1 in intestinal epithelium resulted in altered gastrophysiology and 5-hydroxytryptamine (5-HT) signaling, which was speculated to be responsible for the altered behaviors in NodDKO mice following exposure to stress (*Pusceddu et al., 2019*).

While it is traditionally thought that Nod proteins are cytosolic peptidoglycan (PGN) sensors without recognizable membrane-targeting domains (*Inohara & Nunez, 2003*), several studies have found that Nod1 and Nod2 can be recruited to the plasma membrane and the membranes of endosomes and DCGs (*Bonham & Kagan, 2014; Lu et al., 2019; Travassos et al., 2010; Zhang et al., 2015; Zhang et al., 2019*). Membrane localization of Nod1 and Nod2 has been found to be vital for their function. For example, plasma membrane localization of Nod1 and Nod2 is critical for NF-kB activation in macrophages (*Lu et al., 2019; Nakamura et al., 2014*). Nod1 and Nod2 recruit ATG16L1 onto the plasma membrane to initiate autophagy of bacteria entering the host cell (*Cadwell, 2016; Travassos et al., 2010*). Nod1 and Nod2 are both implicated in processing bacterial outer membrane vesicles through early endosome or autophagosome pathways (*Chu et al., 2016; Travassos et al., 2010*). We previously reported that DCG-localized Nod1 regulates an intracellular membrane trafficking event to promote insulin secretion in pancreatic beta cells (*Zhang et al., 2019*). Therefore, membrane localization of Nod proteins may be essential for fulfilling their diverse functions in different cell contexts.

In this study, we found that Nod1 was highly expressed in adrenal chromaffin cells. We probed for the effect of intestinal microbe-derived Nod1 ligands on hormone secretion in chromaffin cells. Mechanistically, we found that Nod1 localized to DCGs and recruited Rab2a onto DCGs, which subsequently affects CHGA retention in DCGs. Microbe-sensing through Nod1 is required for efficient storage and secretion of catecholamines in adrenal chromaffin cells. Finally, specific deficiency of Nod1 in chromaffin cells impairs epinephrine secretion during immobilization stress in mice. Collectively, our results identify a new microbe-host crosstalk pathway, in which adrenal chromaffin cells directly sense microbial Nod1 ligands released from commensal bacteria by intestinal lysozyme to optimize epinephrine secretion during immobilization stress.

## Results

### Nod1 affects CHGA and epinephrine abundance in adrenal chromaffin cells in a cell-autonomous manner

Immunohistochemical staining in WT mice detected Nod1 expression specifically in the adrenal medulla, but not in the cortex region, while *Nod1* deficiency abolished the medullary staining (*Figure 1A*). In the adrenal gland, *Nod1* mRNA was highly enriched in medulla compared to cortex (*Figure 1—figure supplement 1A*). The medullary *Nod1* mRNA level was comparable to that in spleen, which is known to express Nod1 at a high level (*Figure 1B*), as previously reported (*Inohara et al., 1999*).

**Figure 1.**
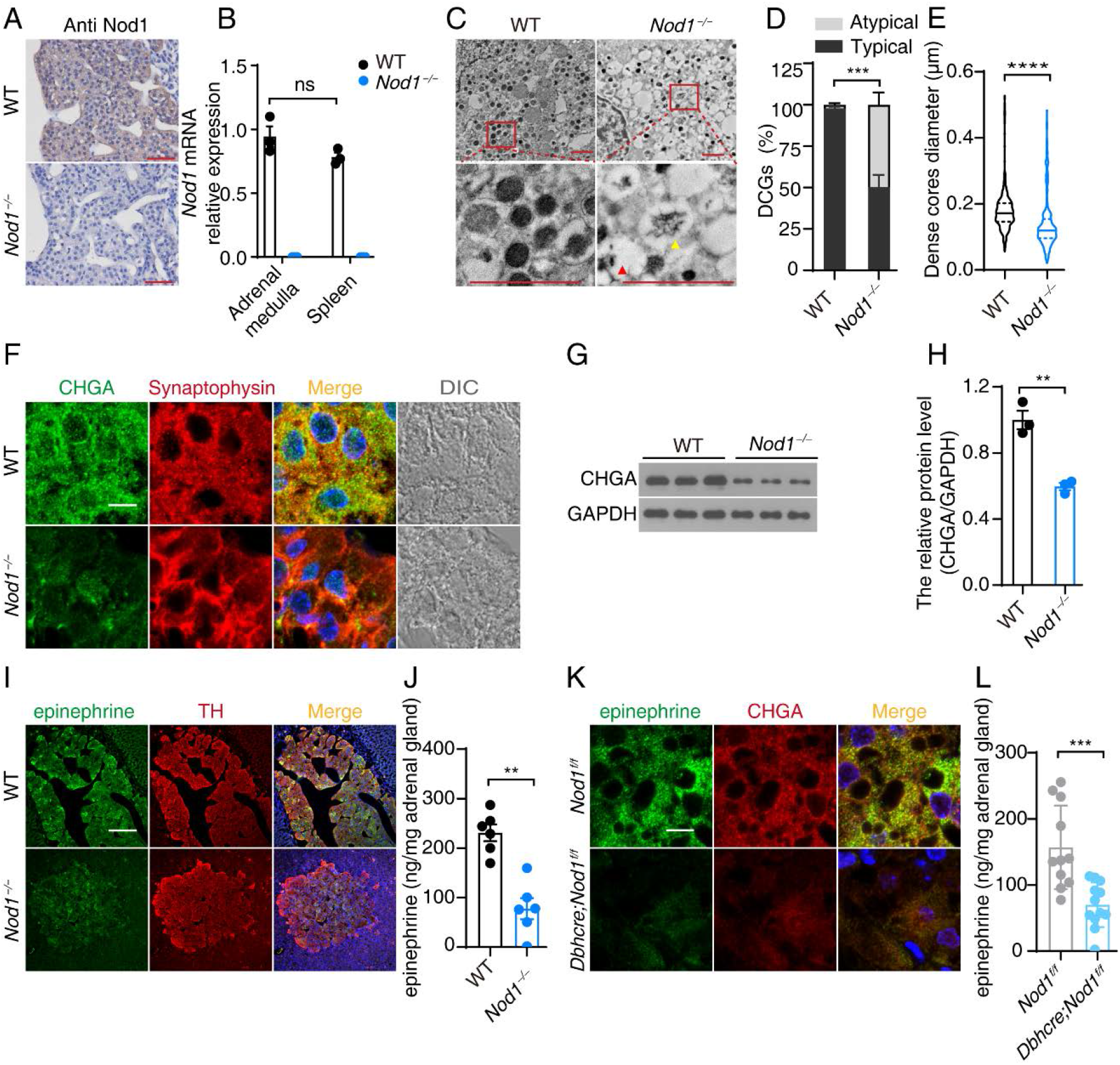
Adrenal chromaffin cells sense microbial signals in a Nod1-dependent manner. (**A**) Immunohistochemical staining (IHC) of Nod1 in paraffin sections of adrenal glands from wild-type (WT) and *Nod1^−/−^* mice. (**B**) Relative *Nod1* mRNA levels in adrenal medulla and spleen determined by qPCR. The *GAPDH* mRNA level was used for normalization. (**C**) Electron microscopy images of adrenal chromaffin cells from WT and *Nod1^−/−^* mice. Boxed areas are shown at higher magnification in the panels immediately below to indicate the atypical dense core granules (DCGs) in *Nod1^−/−^* mice. (**D**) Quantification of DCGs in WT and *Nod1^−/−^* mice from EM images as shown in (**C**). Atypical dense core granules were defined as those with a decreased dense core, as indicated by the yellow triangle in (**C**), or with no noticeable dense core, as indicated by the red triangle in (**C**). A total of approximately 300 DCGs from 2 adrenal gland of each genotype were used for quantification. (**E**) The diameters of DCGs in WT and *Nod1^−/−^* mice. (**F**) Immunostaining and confocal imaging of CHGA (green) and synaptophysin (red) in paraffin sections of adrenal gland from WT and *Nod1^−/−^* mice. (**G, H**) Representative immunoblots (**G**) and quantification (**H**) of CHGA in dissected adrenal medullas from WT and *Nod1^−/−^* mice (n=3). GAPDH was used as loading control. (**I**) Immunostaining and confocal imaging of epinephrine (green) and tyrosine hydroxylase (TH, red) in paraffin sections of adrenal gland from WT and *Nod1^−/−^* mice. (**J**) Adrenal gland epinephrine content in WT and *Nod1^−/−^* mice. (**K**) Immunostaining and confocal imaging of epinephrine (green) and CHGA (red) in paraffin sections of adrenal gland from *Nod1^f/f^* and *Dbh-cre;Nod1^f/f^* mice. (**L**) Adrenal gland epinephrine content in *Nod1^f/f^* and *Dbh-cre;Nod1^f/f^* mice. Scale bars, 10 μm (**F, K**); 1 μm (**C**); 50 μm (**A, I**). Violin plots show median and quartile (**E**). Summary plots show all data points with mean and s.e.m.; each symbol represents an individual mouse (**B, H, J, L**). Statistically significant differences were determined using the Mann-Whitney test (**B, D, J, L**), unpaired student *t test* (**H**) or Kolmogorov-Smirnov test for cumulative distribution (**E**). \*\**p*<0.01, \*\*\**p*<0.001 and \*\*\*\**p*<0.0001. Data (**A-L**) are representative of three independent experiments.

We next sought to understand the biological function of Nod1 in adrenal chromaffin cells. Adrenal chromaffin cells are classic neuroendocrine cells, with DCGs as the hallmark. Transmission electron microscopy (TEM) analysis showed the classic dense core morphology of DCGs in WT chromaffin cells, while a significant portion of DCGs in *Nod1^−/−^* chromaffin cells did not have the typical dense core (*Figure 1C-D*). The diameters of dense cores in *Nod1^−/−^* chromaffin cells tended to be smaller than those in WT chromaffin cells (*Figure 1E*). These data indicate that *Nod1* deficiency led to abnormalities in DCG formation in chromaffin cells.

CHGA plays a key role in the formation of the dense core in DCGs. We probed for CHGA in adrenal medullas of WT and *Nod1^−/−^* mice and found that the CHGA staining signal was significantly reduced in *Nod1^−/−^* medullas compared to WT medullas (*Figure 1F*). Staining of synaptophysin and phogrin, two membrane markers of DCGs, was comparable between WT and *Nod1^−/−^* mice (*Figure 1F* and *Figure 1—figure supplement 1B*). Immunoblotting showed a significantly reduced CHGA protein level in isolated adrenal medullas from *Nod1^−/−^* mice compared to WT mice (*Figure 1G-H*), which confirms our immunofluorescence staining data on tissue sections. *CHGA* mRNA levels were comparable in WT and *Nod1^−/−^* chromaffin cells (*Figure 1—figure supplement 1C*). CHGA is critical for packaging epinephrine in chromaffin cells (*Pasqua et al., 2016*). We used an antibody to probe for epinephrine in adrenal medullas and found that the fluorescence signal of the anti-epinephrine antibody was weaker in medullas from *Nod1^−/−^* mice compared to medullas from WT mice (*Figure 1I*). Tyrosine hydroxylase (TH) was stained to label chromaffin cells (*Figure 1I*). We further quantified the amount of epinephrine in adrenal glands and found that *Nod1* deficiency markedly reduced the abundance of epinephrine (*Figure 1J*).

We wondered whether Nod1 might be involved in affecting the abundance of CHGA and epinephrine in a cell-autonomous manner. To test this possibility, we specifically deleted Nod1 in adrenergic and noradrenergic cells by generating *Dbh-cre;Nod1^f/f^* mice, in which loxP-flanked *Nod1* alleles (*Nod1^f/f^*) were deleted by Cre recombinase driven by the dopamine beta hydroxylase (Dbh) promoter). In *Dbh-cre;Nod1^f/f^* mice, the *Nod1* mRNA level was markedly reduced in adrenal gland, but not in other tissues, such as spleen or lung (*Figure 1—figure supplement 1D*). Specific deletion of Nod1 in chromaffin cells led to marked reduction in staining of epinephrine and CHGA (*Figure 1K*). Accordingly, adrenal glands from *Dbh-cre;Nod1^f/f^* mice contained less epinephrine than control *Nod1^f/f^* mice (*Figure 1L*). Thus, deficiency of Nod1 led to reduced levels of CHGA and epinephrine in DCGs in a cell-autonomous manner.

### Microbial Nod1 ligand regulates the amount of CHGA and epinephrine in DCGs

Nod1 is a cytosolic receptor which has been implicated in recognizing peptidoglycan ligands from pathogens as well as intestinal commensals (*Chamaillard et al., 2003*). We suspected that Nod1 ligand from intestinal microbes might be involved in regulating Nod1 in adrenal chromaffin cells. Treatment with antibiotics has been used to deplete microbial ligands in the circulation (*Arentsen et al., 2017; Bird, 2010; Clarke et al., 2010; Hergott et al., 2016*). We previously reported that the presence of circulating Nod1 ligand depends on lysozyme in the intestinal lumen, and deficiency of Lyz1 in mice leads to depletion of microbial Nod1 ligands (*Zhang et al., 2019*). To determine the effect of intestinal Nod1 ligand on Nod1 in chromaffin cells, we used mice treated a cocktail of antibiotics (ABX) and *Lyz1^−/−^* mice. We compared Nod1 staining in WT mice, ABX mice and *Lyz1^−/−^* mice. We found that Nod1 localized to CHGA^+^ DCGs in specific pathogen-free (SPF) WT mice, but the staining of Nod1 on CHGA^+^ DCGs was very weak in germ-free (GF) mice, which lack gut microbiota (*Figure 2A*). Depletion of commensal bacteria by antibiotic cocktail treatment or depletion of Lyz1 diminished the staining of Nod1 on CHGA^+^ DCGs (*Figure 2A*). mRNA quantification and IHC of adrenal medullas showed that ABX treatment or loss of Lyz1 did not affect the protein expression of Nod1 (*Figure 2—figure supplement 1A-B*). This indicates that the diminished staining of Nod1 in chromaffin cells in ABX or *Lyz1^−/−^* mice was due to reduced recruitment of Nod1 onto DCGs.

**Figure 2.**
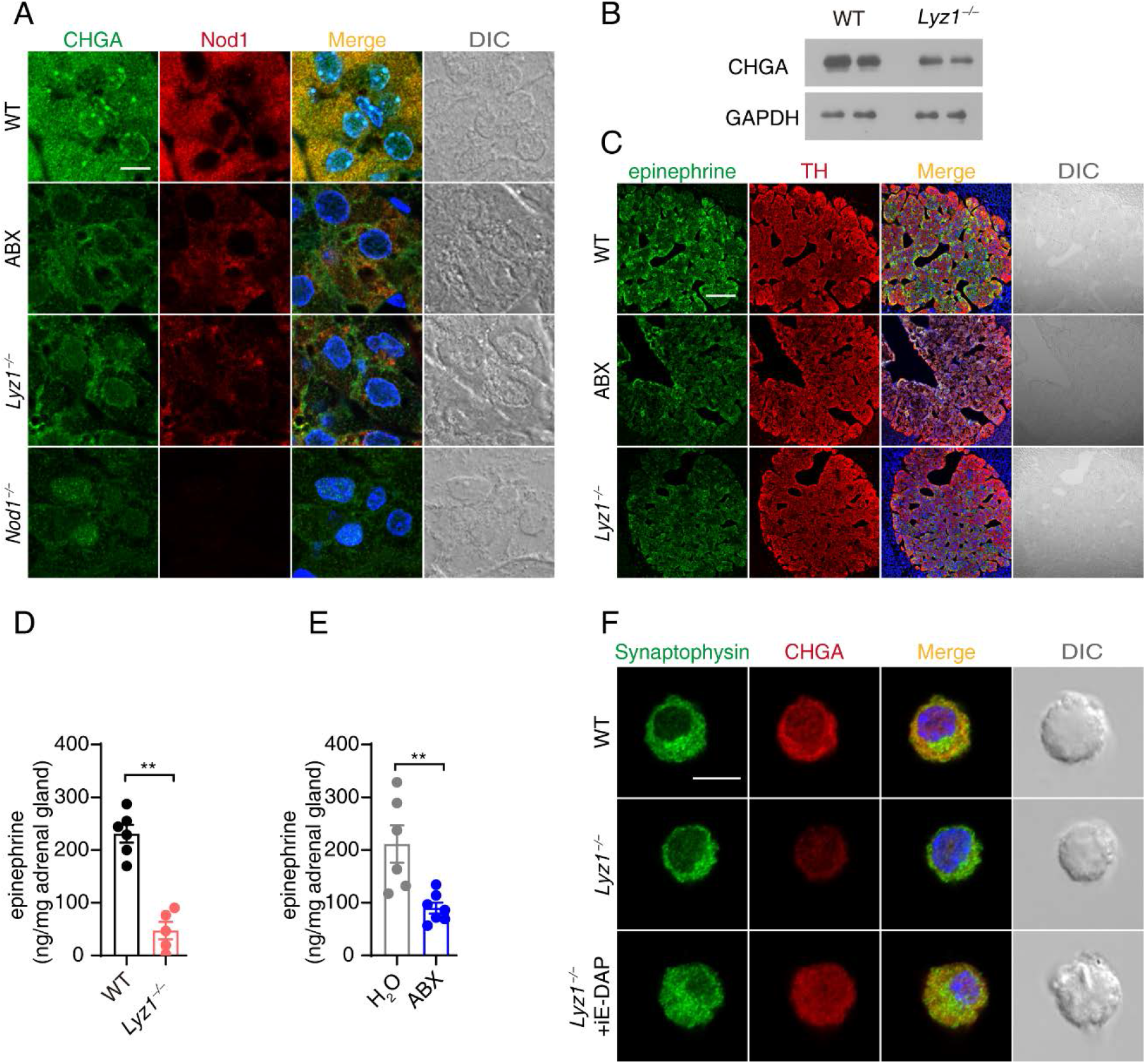
Microbial Nod1 ligand regulates the amount of CHGA and epinephrine in DCGs. (**A**) Immunostaining and confocal imaging of CHGA (green) and Nod1 (red) in paraffin sections of adrenal gland from WT, ABX, *Lyz1^−/−^* and *Nod1^−/−^* mice. (**B**) Representative immunoblots of CHGA in the dissected adrenal medullas from WT and *Lyz1^−/−^* mice. GAPDH was used as loading control. (**C**) Immunostaining and confocal imaging of epinephrine (green) and TH (red) in paraffin sections of adrenal gland from WT, ABX and *Lyz1^−/−^* mice. (**D**) The epinephrine content in adrenal glands from WT (n=6) and *Lyz1^−/−^* (n=5) mice. (**E**) The epinephrine content in adrenal glands from mice treated with H_2_O (vehicle, n=6) or antibiotic cocktail (ABX, n=7). (**F**) Immunostaining and confocal imaging of synaptophysin (green) and CHGA (red) in primary cultured chromaffin cells from WT and *Lyz1^−/−^* mice (treated with iE-DAP or PBS). Scale bars, 10 μm (**A, F**); 50 μm (**C**). Statistically significant differences were determined using the Mann-Whitney test (**D, E**). \*\**p*<0.01 and \*\*\*\**p*<0.0001. Each symbol represents an individual mouse. Mean and s.e.m. are indicated (**D, E**). Data (**A-F**) are representative of three independent experiments.

Similar to the observation in *Nod1^−/−^* mice, CHGA staining was greatly diminished in ABX or *Lyz1^−/−^* mice (*Figure 2A* and *Figure 2—figure supplement 1C*). In comparison, localization of Nod1 on Golgi was not affected in ABX mice or *Lyz1^−/−^* mice compared to WT mice (*Figure 2—figure supplement 1D*). Significantly less CHGA was detected in isolated adrenal glands of *Lyz1^−/−^* mice compared to WT mice (*Figure 2B*). Immunofluorescence staining detected marked reduction of epinephrine in TH^+^ cells in adrenal gland in ABX mice or *Lyz1^−/−^* mice compared to WT mice (*Figure 2C*). Accordingly, the levels of epinephrine and CHGA were both reduced in adrenal gland in ABX mice and *Lyz1^−/−^* mice compared to WT mice (*Figure 2—figure supplement 1E*). The reduction in epinephrine in adrenal gland in ABX mice and *Lyz1^−/−^* mice was confirmed by ELISA quantification (*Figure 2D-E*). Furthermore, the reduction in CHGA in chromaffin cells from *Lyz1^−/−^* mice was observed in cultured primary chromaffin cells, and treatment with the cognate Nod1 ligand, iE-DAP, restored CHGA in chromaffin cells from *Lyz1^−/−^* mice (*Figure 2F*). This indicates that Nod1 ligand is required to maintain the CHGA level in chromaffin cells.

### Nod1 recruits Rab2a onto DCGs to regulate CHGA retention in DCGs

Rabs, a family of small GTPases, are master regulators of intracellular membrane trafficking. Rab2a is known to be a Golgi-resident GTPase, governing intra-Golgi trafficking (*Ailion et al., 2014; Buffa, Fuchs, Pietropaolo, Barr, & Solimena, 2008*). In addition, Rab2a localizes to the membranes of lysosome-like organelles, such as DCGs, melanosomes and early endosomes. Studies have shown that Rab2a regulates cargo sorting events in neurons in *C. elegans* and intestinal Paneth cells (*Edwards et al., 2009; Sumakovic et al., 2009; Zhang et al., 2015*). In *C. elegans*, Rab2 null mutation impaired the maturation of the dense core of DCGs, and DCG cargos were mistrafficked to early endosomes for degradation (*Edwards et al., 2009; Sumakovic et al., 2009*). In intestinal Paneth cells, Rab2a knockdown led to mistargeting of lysozyme to lysosomes for degradation instead of storage in DCGs (*Zhang et al., 2015*). Thus, we suspected that Rab2a might act downstream of Nod1 in regulating CHGA storage in DCGs. We performed immunostaining of Rab2a and Nod1 in adrenal medulla. Rab2a and Nod1 largely colocalized in WT chromaffin cells; however, colocalization of Rab2a and Nod1 was largely lost in chromaffin cells from ABX or *Lyz1^−/−^* mice (*Figure 3A*). In contrast, the colocalization of Rab2a with Nod1 around the nucleus was not affected in chromaffin cells from ABX or *Lyz1^−/−^* mice (*Figure 3A*). Nod1 localizes to DCGs and the perinuclear Golgi (*Figure 2A* and *Figure 2—figure supplement 1D*); therefore, we suspect that the remaining colocalization of Nod1 and Rab2a in the perinuclear region was Golgi localization, which remained undisturbed in chromaffin cells from ABX or *Lyz1^−/−^* mice. To test that possibility that Nod1 specifcially affects Rab2a localization onto DCGs, we performed immunostaining of Rab2a together with GM130, a Gogli marker. We found that, in addition to localizing to Golgi marked by GM130, Rab2a staining was also observed on puncta throughout in the cytosol in the control *Nod1^f/f^* mice (*Figure 3B*). However, specific depletion of Nod1 in chromaffin cells in *Dbh-cre;Nod1^f/f^* mice resulted in the loss of Rab2a staining on puncta in the cytosol (*Figure 3B*). In comparison, the localization of Rab2a on Gogli was not affected in chromaffin cells in *Dbh-cre;Nod1^f/f^* mice (*Figure 3B*).

**Figure 3.**
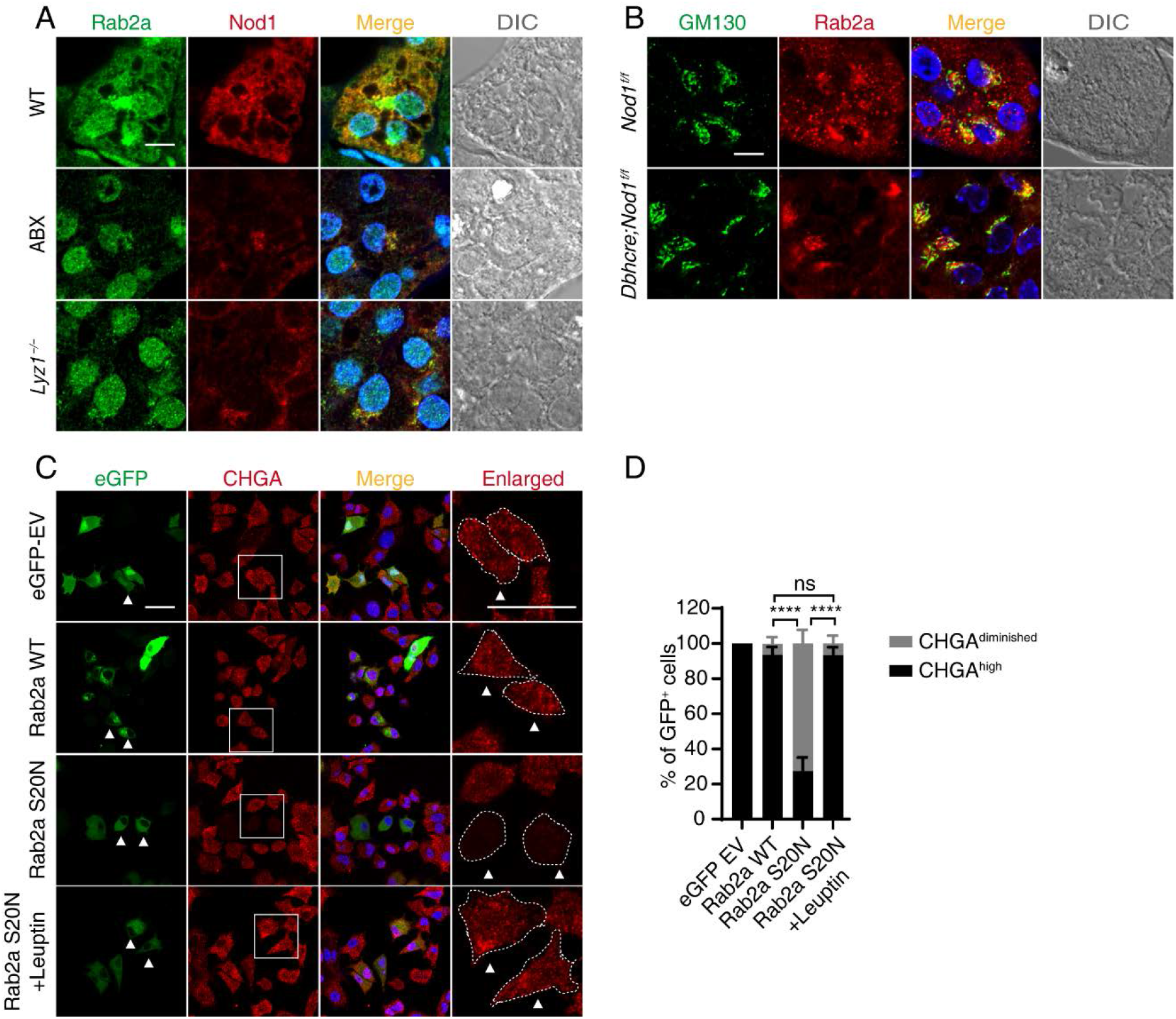
Nod1 recruits Rab2a onto DCGs to regulate CHGA retention in DCGs. (**A**) Immunostaining and confocal imaging of Rab2a (green) and Nod1 (red) in paraffin sections of adrenal gland from WT, ABX and *Lyz1^−/−^* mice. (**B**) Immunostaining and confocal imaging of GM130 (green) and Rab2a (red) in paraffin sections of adrenal gland from *Nod1^f/f^* and *Dbh-cre;Nod1^f/f^* mice. (**C**) PC-12 cells were transfected with eGFP empty vector (EV), wild-type Rab2a tagged with eGFP, or a dominant-negative Rab2a-S20N mutant tagged with eGFP. Cells were immunostained with anti-CHGA (red) antibody. Cells in the boxed regions are highly magnified on the right. White dashed line outlines the designated cell boundry. Arrowheads indicate representative GFP^+^ cells. (**D**) Quantification of transfected (GFP^+^) CHGA^diminished^ cells and CHGA^high^ cells from (**C**). Cells were counted from 8-10 fields of view with at least 140 GFP^+^ cells quantified for each sample. Scale bars, 10 μm in (**A**, **B**), 30 μm in (**C**). Statistically significant differences were determined using one-way ANOVA followed by Tukey’s post hoc tests (**D**). \*\*\*\**p*<0.0001. ns indicates no significant difference. Summary plot shows means and s.e.m. (**D**). Data (**A-D**) are representative of three independent experiments.

To determine whether Rab2a affects CHGA retention in DCGs, we determined the effect of Rab2a on CHGA in PC-12 cells, derived from a pheochromocytoma of the rat adrenal medulla (*Greene & Tischler, 1976*). To perturb the activity of Rab2a, we created a Rab2a mutant, S20N, which was previously determined as a dominant negative mutant (*Tisdale, Bourne, Khosravi-Far, Der, & Balch, 1992*). Expression of Rab2a-WT did not alter the level of CHGA in transfected cells; however, expression of the S20N mutant resulted in significant reduction of CHGA (*Figure 3C-D*). It was previously shown that loss of Rab2a mistargeted DCG cargos for degradation in lysosomes (*Edwards et al., 2009; Sumakovic et al., 2009; Zhang et al., 2015*). We treated PC-12 cells expressing Rab2a S20N with a lysosome inhibitor, leupeptin. The leupeptin treatment effectively restored CHGA in cells expressing the Rab2a S20N mutant (*Figure 3C-D*). Thus, Rab2a activity is involved in regulating the level of CHGA in DCGs.

### Reduced catecholamine secretion in chromaffin cells deficient in Nod1 or Nod1 ligand

We next decided to determine whether reduced storage of CHGA and epinephrine could lead to reduced hormone secretion. Single-cell amperometry measures the numbers of released catecholamine molecules during every individual exocytosis event (*Garcia, Garcia-De-Diego, Gandia, Borges, & Garcia-Sancho, 2006; Wightman et al., 1991*). Basically, catecholamines from each vesicle are captured and oxidized on a carbon electrode, resulting in a current (or amperometric) peak for each vesicle that bursts (*Figure 4A*). Each validated current peak presents a fusion event, and the number of catecholamine molecule from each fusion event is calculated from the integration of each single current event (*Garcia et al., 2006; Wightman et al., 1991*). Single-cell amperometry does not distinguish different kinds of catecholamines, but it can accurately quantify the number of molecules released and the kinetic parameters of each vesicle (*Dunevall et al., 2015*). Single-cell amperometry was employed to record each K^+^-stimulated fusion event between secretory vesicles and the plasma membrane on cultured primary chromaffin cells from WT, *Nod1^−/−^* and *Lyz1^−/−^* mice (*Figure 4B*). The peak current of individual amperometric spike events was distributed toward reduced current in *Nod1^−/−^* and *Lyz1^−/−^* chromaffin cells (*Figure 4C*), and the means of peak current were reduced in *Nod1^−/−^* and *Lyz1^−/−^* chromaffin cells compared with WT cells (*Figure 4D*). The catecholamine molecule numbers from each fusion event was calculated. The distribution of catecholamine molecule numbers, together with the mean numbers of secreted catecholamine molecules per event, indicated that fewer catecholamine molecules were secreted from *Nod1^−/−^* and *Lyz1^−/−^* chromaffin cells compared with WT cells (*Figure 4E-F*). Taken together, these data indicate that the deficiency of Nod1 or Lyz1 leads to impaired catecholamine secretion in DCGs.

**Figure 4.**
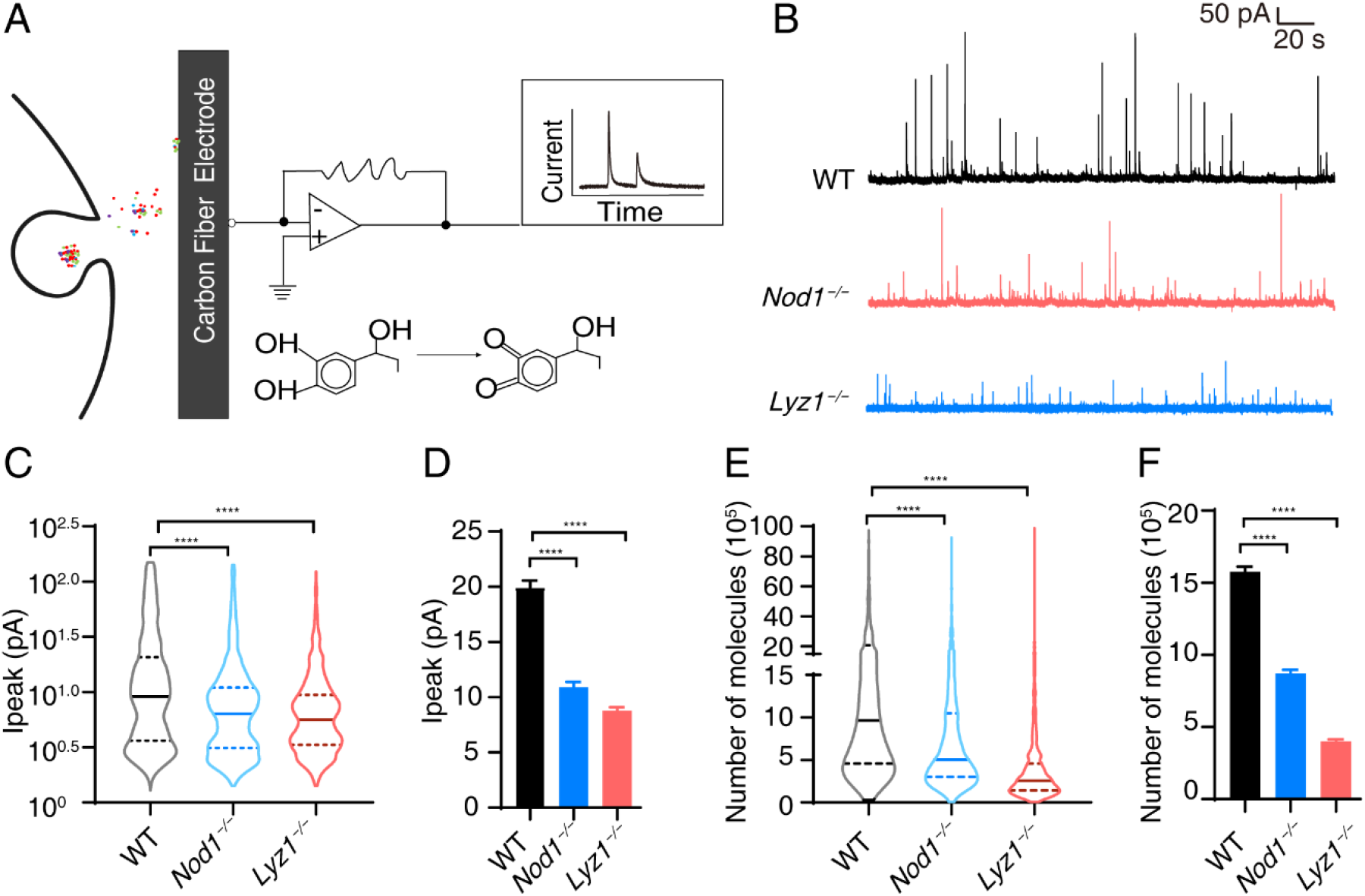
Amperometry recordings reveal a significant decrease in the number of catecholamine molecules released by individual vesicles in *Nod1^−/−^* and *Lyz1^−/−^* adrenal chromaffin cells. (**A**) Schematic diagram of the amperometry recording experiment. (**B**) Representative amperometric traces from WT, *Nod1^−/−^* and *Lyz1^−/−^* chromaffin cells. (**C**) Violin plot of peak current of each amperometric spike event from WT (n=2031 events), *Nod1^−/−^* (n=1243 events) and *Lyz1^−/−^* (n=1317 events) chromaffin cells. (**D**) Means of peak current from (**C**) were plotted with s.e.m. (**E**) Violin plot of number of molecules in each amperometric spike event from WT (n=2172 events), *Nod1^−/−^* (n=1329 events) and *Lyz1^−/−^* (n=1341 events) chromaffin cells. (**F**) Means of number of molecules from (**E**) were plotted with s.e.m. Statistically significant differences were determined using one-way ANOVA followed by Tukey’s post hoc tests (**C-F**). \*\**p*<0.01.

Single-cell amperometry does not distinguish epinephrine and norepinephrine from other monoamines, including dopamine (*Robinson, Venton, Heien, & Wightman, 2003*). Specific probes have been developed to monitor catecholamine dynamics in a real-time manner (*Muller, Joseph, Slesinger, & Kleinfeld, 2014; Wang, Jing, & Li, 2018*). A GPCR activation-based NE (GRAB_NE_) sensor, based on an α-2 adrenergic receptor, has been used for *in vivo* and *in vitro* measurement of epinephrine or norepinephrine dynamics in mice (*Feng et al., 2019*). The GRAB_NE_ sensor has circularly permutated eGFP (cp eGFP) inserted between the fifth and sixth transmembrane domains of the α-2 adrenergic receptor. When epinephrine or norepinephrine binds to the extracellular domain of the GRAB_NE_ sensor, a conformational change occurs between the fifth and sixth transmembrane domains to activate the green fluorescence of cp eGFP. Thus, when GRAB_NE_ sensor is expressed on the cell surface, its fluorescence reflects the level of epinephrine/norepinephrine in the extracellular environment (*Feng et al., 2019*). Adrenal chromaffin cells mostly secrete epinephrine, so we employed the GRAB_NE_ method to evaluate whether Nod1 or Nod1 ligand could affect the secretion of epinephrine (and, to a lesser extent, norepinephrine). Chromaffin cells from WT, *Nod1^−/−^* or *Lyz1^−/−^* mice were isolated and transfected with GRAB_NE_ sensor. As control, RFP was co-transfected to provide continuous fluorescence in the cytoplasm, as previously described (*Feng et al., 2019*). The green fluorescence on the plasma membrane in WT cells increased rapidly when cells were exposed to a high concentration of K^+^, and then gradually decreased and returned to the ground state as time elapsed (*Figure 5A*). RFP in the cytoplasm remained constant during the course of K^+^ stimulation (*Figure 5—figure supplement 1*). In comparison, K^+^ stimulation did not prompt a strong enhancement in the green fluorescence on the plasma membrane of the transfected *Nod1^−/−^* or *Lyz1^−/−^* chromaffin cells (*Figure 5A*). The fluorescence intensity changes of GRAB_NE_ on the plasma membrane and of RFP were recorded and quantified. Loss of *Nod1* or *Lyz1* in mice resulted in a weaker increase in the fluorescence intensity of the GRAB_NE_ sensor, compared to WT mice (*Figure 5B*). The maximum fluorescence intensity change of GRAB_NE_ was also reduced in chromaffin cells from *Nod1^−/−^* or *Lyz1^−/−^* mice, compared to cells from WT mice (*Figure 5C*). These data indicate that less epinephrine (norepinephrine) was secreted by cells from *Nod1^−/−^* or *Lyz1^−/−^* mice. We further supplemented cultured primary chromaffin cells from *Lyz1^−/−^* mice with iE-DAP. Such treatment restored the K^+^-stimulated fluorescence augmentation of the GRAB_NE_ sensor (*Figure 5B-C*), which indicates that the absence of Nod1 ligand was responsible for the decreased secretion of epinephrine by cells from *Lyz1^−/−^* mice. In addition, chromaffin cells isolated from *Dbh-cre;Nod1^f/f^* mice also displayed a similar reduction in epinephrine (norepinephrine) secretion (*Figure 5D-F*), further supporting the idea that Nod1 plays a cell-autonomous role in regulating secretion in chromaffin cells.

**Figure 5.**
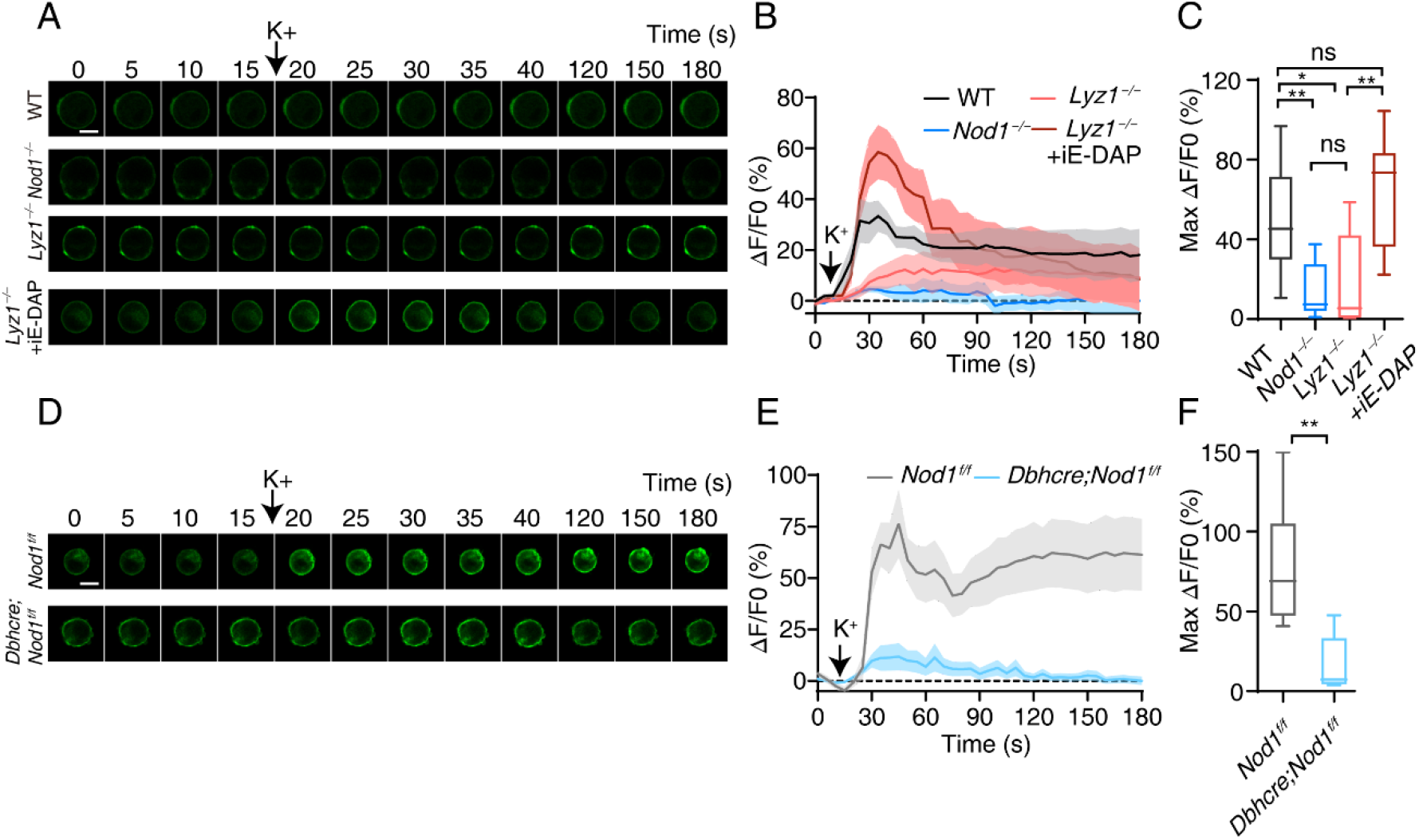
Reduced catecholamine secretion in chromaffin cells deficient in Nod1 or Nod1 ligand. (**A**) Representative images of green fluorescence of GRAB_NE_ in transfected chromaffin cells from WT, *Nod1^−/−^* and *Lyz1^−/−^* (pretreated with iE-DAP for 6 h or PBS) during the course of treatment with K^+^ solution (90 mM) to trigger secretion. (**B**) The recorded fluorescence changes of GRAB_NE_ over the time course of the experiment (ΔF/F0) (n=8-12 cells from 5-8 mice). (**C**) Box plot of the maximum peak of ΔF/F0 from (**B**) with median and interquartile range indicated. (**D**) Representative images of green fluorescence of GRAB_NE_ in transfected chromaffin cells from *Nod1^f/f^* and *Dbh-cre;Nod1^f/f^* mice during the course of treatment with K^+^ solution (90 mM). (**E**) The recorded fluorescence changes of GRAB_NE_ over the time course of the expriment (ΔF/F0) (n=6-10 cells from 5-8 mice). (**F**) Box plot of the maximum peak of ΔF/F0 from (**E**). The line connects means of fluorescence changes over the time course with ribbons indicating ± s.e.m. (**B, E**). Scale bars, 10 μm (**A, D**). Statistically significant differences were determined using one-way ANOVA followed by Tukey’s post hoc tests (**C**) and Student’s *t*-test (**F**). \**p*<0.05 \*\**p*<0.01, \*\*\**p*<0.001 and \*\*\*\**p*<0.0001.

### Deficiency of Nod1 ligand impairs the stress response

We wanted to determine the physiological function of Nod1 sensing in adrenal chromaffin cells. We employed a mouse immobilization model, which has been widely used to evaluate the adrenal medullary response during stress. The elevated epinephrine, together with other stress hormones, induces a range of physiological responses in preparation for the fight-or-flight response, including increased blood sugar level, glycogen mobilization in the liver, *etc*. (*Buira et al., 2004; Fernandez et al., 2000; Kang, Sim, Park, Sharma, & Suh, 2015; Park et al., 2016*). We performed the immobilization with WT and *Lyz1^−/−^* mice. WT and *Lyz1^−/−^* mice had comparable levels of circulating epinephrine under basal (non-stressed) conditions (*Figure 6A*). When the mice were immobilized for one hour, the level of circulating epinephrine markedly increased in WT mice, but the increase was blunted in *Lyz1^−/−^* mice (*Figure 6A*). Prior studies have found that immobilization raises the blood sugar level (*Kang et al., 2015; Park et al., 2016*). We found that the surge in blood sugar was clearly detected in both *Lyz1^−/−^* and WT mice after restraint (*Figure 6B*).

**Figure 6.**
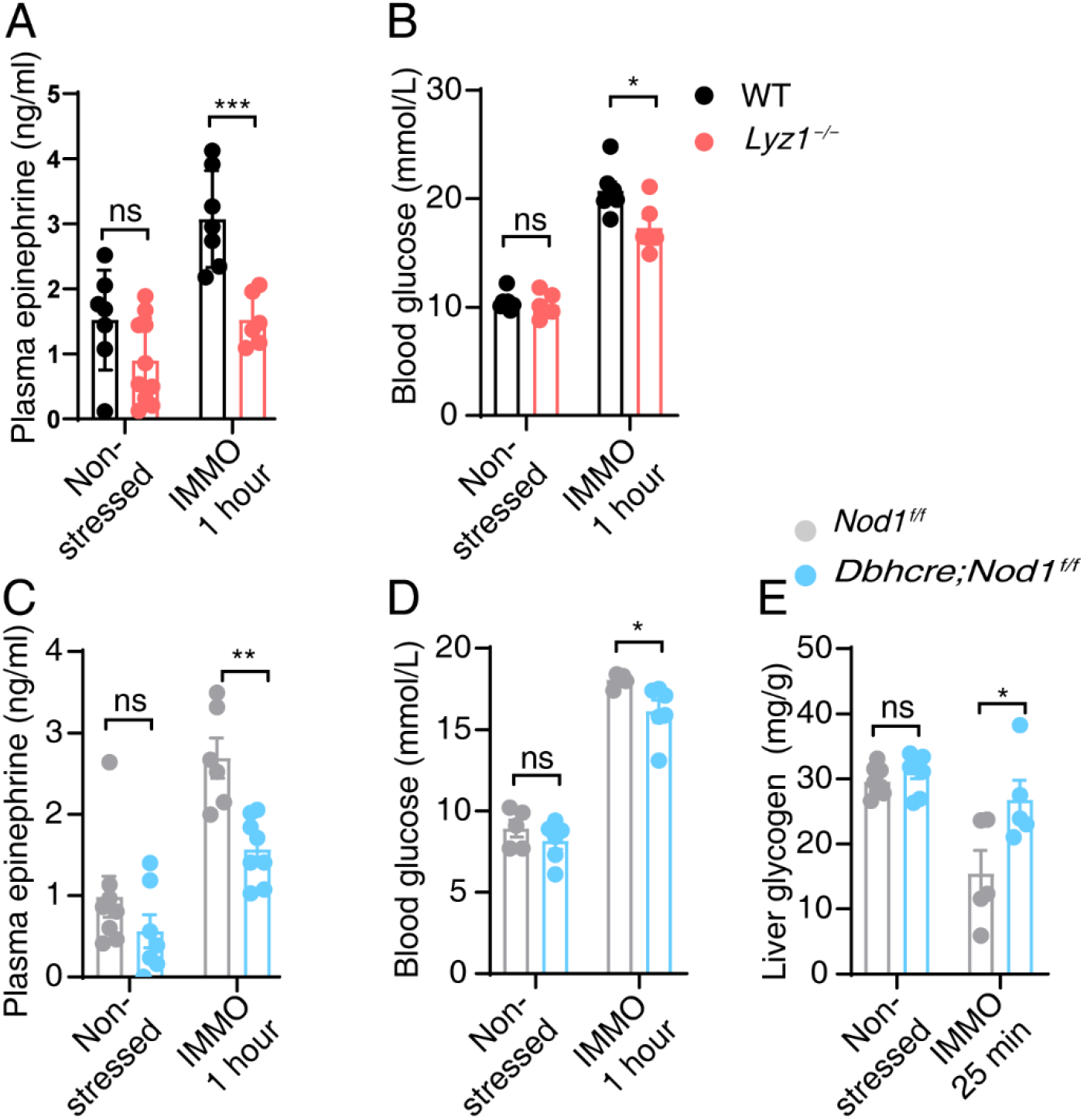
Deficiency of Nod1 ligand impairs epinephrine response. Deficiency of Nod1 ligand impairs epinephrine response. (**A, C**) Plasma level of epinephrine in the steady state and following stress induction by immobilization (IMMO; 1 h) in WT, *Lyz1^−/−^, Nod1^f/f^* and *Dbh-cre;Nod1^f/f^* mice. (**B, D**) Concentration of blood glucose in the steady state and following immobilization (IMMO; 1 h) in WT, *Lyz1^−/−^, Nod1^f/f^* and *Dbh-cre;Nod1^f/f^* mice. (**E**) Liver glycogen content under steady-state conditions or after 25 min IMMO in *Nod1^f/f^* and *Dbh-cre;Nod1^f/f^* mice. Statistically significant differences were determined using one-way ANOVA followed by Tukey’s post hoc tests (**A-E**). ns, indicates no significant difference (*p*>0.05). \**p*<0.05, \*\**p*<0.01 and \*\*\**p*<0.001. Summary plots (**A-E**) show mean with s.e.m. Each symbol represents an individual mouse.

To further determine the cell-autonomous function of Nod1 in adrenal medulla, we evaluated control and *Dbh-cre;Nod1^f/f^* mice under immobilization. Control and *Dbh-cre;Nod1^f/f^* mice had a comparable level of circulating epinephrine (*Figure 6C*). Immobilization resulted in a dramatic increase in the circulating epinephrine level in control mice, while the increase in epinephrine level was less pronounced in *Dbh-cre;Nod1^f/f^* mice (*Figure 6C*). The increase in blood sugar was also blunted in *Dbh-cre;Nod1^f/f^* mice compared to control mice (*Figure 6D*). Furthermore, the stressed-induced depletion of glycogen store in liver in *Dbh-cre;Nod1^f/f^* mice was milder compared to control mice (*Figure 6E*). Thus, our data demonstrate that Nod1 sensing in adrenal medulla is involved in epinephrine secretion during immobilization stress.

## Discussion

Here we report that commensal bacteria are sensed directly in adrenal chromaffin cells to modulate epinephrine output during immobilization stress. Upon sensing its ligand, Nod1 recruits Rab2a onto DCGs, which assists the condensation of CHGA in DCGs. Defects in this pathway lead to reduced epinephrine secretion in mouse. Thus, our study unveils a physiological role of Nod1-mediated crosstalk between intestine and adrenal medulla to promote the fight-or-flight response.

Our study investigated the effect of Nod1 on epinephrine secretion in the adrenal medullary response. In our study, adrenal chromaffin cells expressed Nod1 at a high level, and depletion of Nod1 in adrenal chromaffin cells led to a dramatic reduction in CHGA and epinephrine. The effects on CHGA and epinephrine were observed in cultured primary chromaffin cells, thus supporting a cell-autonomous role of Nod1 in chromaffin cells. Supplementing cultured primary chromaffin cells from *Lyz1^−/−^* mice with Nod1 ligand *in vitro* restored epinephrine secretion, which argues for a direct effect of microbial ligands on chromaffin cells. We employed conditional Nod1 knockout mice to further establish the cell-autonomous role of Nod1 in chromaffin cells. In *Dbh-cre;Nod1^f/f^* mice, Nod1 was depleted in adrenal chromaffin cells and Dbh-expressing (Dbh^+^) neurons. Adrenal chromaffin cells are the main source of epinephrine during stress (*Axelrod & Weinshilboum, 1972*). We observed a major reduction in the level of circulating epinephrine in *Dbh-cre; Nod1^f/f^* mice compared to control mice. Thus, we propose that Nod1 sensing in adrenal chromaffin cells regulates epinephrine secretion and consequently plays a critical role in the stress response. Of course, we cannot exclude the possibility that Nod1 may play roles in other catecholaminergic Dbh^+^ neurons during the stress response. It remains to be determined whether Nod1 expressed in cells other than intestinal epithelium and adrenal chromaffin cells may regulate other aspects of physiological processes involved in the stress response.

We found that loss of microbial Nod1 ligand results in impaired storage and secretion of epinephrine in DCGs. Mechanistically, we found that *Nod1^−/−^*, *Lyz1^−/−^*, and *Dbh-cre;Nod1^f/f^* mice contained lower levels of CHGA and epinephrine. CHGA has been shown to regulate catecholamine-containing dense-core chromaffin granule biogenesis in the adrenal gland (*Díaz-Vera et al., 2012; Kim et al., 2001; Pasqua et al., 2016*). We attribute the reduced storage of epinephrine to the reduced amount of CHGA. The previous finding found that the absence of gut microbial colonization selectively impaired catecholamine response to hypoglycemic stress in mice, and transcriptome in adrenal glands was altered (*Giri et al., 2019*). It is possible that Nod1 regulated storage of epinephrine may also contribute to impaired catecholamine response to hypoglycemic stress.

CHGA plays a key role in epinephrine storage. Our results found that mRNA levels of CHGA was not affected while protein levels of CHGA were reduced in adrenal medullas in *Nod1^−/−^*, *Lyz1^−/−^*, and *Dbh-cre;Nod1^f/f^* mice. Thus, we did not perform whole-genome transcriptome profiling, and instead, we focused our investigation on membrane trafficking event of CHGA. We observed reduced recruitment of Rab2a onto DCGs in *Nod1^−/−^*, *Lyz1^−/−^*, and *Dbh-cre;Nod1^f/f^* mice. Overexpressing a dominant negative mutant of Rab2a reduced the level of CHGA in PC-12 cells, which could be restored by inhibiting lysosomes. We propose a scenario where a reduced level of Nod1 or Nod1 ligand impairs Rab2a recruitment onto DCG membranes, which may cause mistargeting of a significant proportion of CHGA to lysosomes for degradation. The involvement of Rab2a in DCG cargo sorting has been previously reported (*Edwards et al., 2009; Sugawara, Kano, & Murata, 2014; Sumakovic et al., 2009; Zhang et al., 2015*). Loss of Rab2a in neurons in *C. elegans* led to specific lysosomal degradation of a neuropeptide that is normally located in DCGs (*Edwards et al., 2009; Sumakovic et al., 2009*). Knockdown of Rab2a led to lysosomal degradation of lysozyme in intestinal Paneth cells (*Zhang et al., 2015*). Besides DCGs, Rab2a also functions at various steps of intracellular membrane trafficking, including ER-Golgi transport, autophagy and autolysosome formation, and endosome-lysosome fusion. Indeed, our results show that Nod1 also localizes to the Golgi. However, Rab2a localization around the Golgi was not affected by loss of Nod1. Therefore, by unknown mechanisms, Nod1 together with Nod1 ligand is required to recruit Rab2a onto DCGs. We suspect some other factors downstream of Nod1 are involved in recruiting Rab2a onto DCGs, which warrants further investigation.

It was previously reported that Nod1 together with Rip2 recruited Rab1a onto insulin granules in islet beta cells to regulate an anterograde transport event (*Zhang et al., 2019*). We found no difference in Rab1a localization on DCGs in chromaffin cells in WT and *Nod1^−/−^* mice (data not shown). It is intriguing that Nod1 recruits different Rab proteins onto vesicles in different cell types. Additional factors may be involved in recruiting specific downstream Rab proteins in different cell types.

In summary, Nod1 ligand from intestinal microbes directly modulates the storage of CHGA and epinephrine in DCGs in adrenal chromaffin cells, which optimizes epinephrine secretion under stress conditions. Thus, our study further exemplifies Nod1 as a key signaling molecule in the microbiota-intestine-brain axis.

## Experimental model and subject details

### Mice

*Lyz1^−/−^, Nod1^−/−^* and *Nod1^f/f^* mice (C57BL/6 background) were generated using the CRISPR/Cas9 method as described previously (*Zhang et al., 2019*). The *Dbh-Cre* (Tg(Dbh-cre)KH212Gsat/Mmucd) mice were a gift from Professor Chen Zhan (NIBS) and were originally from the Mutant Mouse Resource & Research Center (MMRRC) (*Gerfen, Paletzki, & Heintz, 2013*). The *Dbh-cre* mice were further crossed with *Nod1^f/f^* to generate conditional knockout (*Dbh-cre;Nod1^f/f^)* mice. 2-4-month-old gender-matched mice (both male and female) were used in this study unless noted otherwise. All mice were bred at 3-6 animals per cage in a 12/12 h light-dark cycle with *ad libitum* access to food and water at a controlled temperature (23°C ± 2°C). All specific pathogen-free (SPF) mice were bred and housed in an AAALAC-accredited barrier facility in Tsinghua University. All procedures were approved by the Committee of Institutional Animal Care and Research, Tsinghua University.

## Method details

### Plasmids

Human Rab2a cDNA was reverse-transcribed from mRNA prepared from HEK293T cells and inserted into peGFP-C1 vector. The dominant-negative (S20N) Rab2a mutant was generated by PCR-mediated mutagenesis. Sequences were confirmed by Sanger sequencing.

### Cell culture and immunofluorescence

PC-12 cells were cultured in RPMI-1640 medium supplemented with 10% fetal bovine serum and 1% Pen/Strep. The cells were cultured at 37°C in a humidified atmosphere containing 5% CO_2_. For immunofluorescence experiments, PC-12 cells were transfected with 3 μg plasmid and 9 μl Lipofectamine 2000 in 300 μl Opti-MEM media for 6 hours. Cells were incubated at 37°C for 24 hours prior to testing for eGFP, Rab2a WT and Rab2a S20N expression. Cultured PC-12 cells were treated with leupeptin (100 mM) or were mock treated for 24 h in growth medium before being processed for CHGA immunostaining.

### Isolation and culture of mouse adrenal chromaffin cells

Adrenal glands from 2-4 months old WT, *Lyz1^−/−^, Nod1^−/−^, Dbh-cre;Nod1^f/f^* and *Nod1^f/f^* mice were used in each culture. Animals were anesthetized with CO_2_, and the glands were isolated following the procedures described by Aaron Kolski-Andreaco *et al*. (*Kolski-Andreaco, Cai, Currle, Chandy, & Chow, 2007*). Glands were placed into a dish containing oxygenated Locke’s buffer (NaCl 140 mM, KCl 5 mM, HEPES 10 mM, glucose 10 mM, MgCl_2_ 1.2 mM, CaCl_2_ 2.2 mM; pH7.35, osmolarity between 295 and 300 mOsm.) on ice, and cortexes were removed. Adrenal medullas were digested for 25 minutes twice in DMEM with papain (40 unit/ml), 1 mM CaCl_2_ and 0.5 mM EDTA at 37°C. Subsequently, the medullas were disrupted with a yellow tip in complete medium with DMEM, 5% fetal calf serum,10% horse serum, and 5 μl/ml penicillin and streptomycin. Cells were pelleted for 8 minutes at 200 g. Pellets were resuspended in complete medium and plated on poly-L-lysine pretreated coverslips or in imaging chambers. Primary adrenal chromaffin cells were then kept in an incubator at 37°C with 5% CO_2_.

### Amperometric recording and solutions

Isolated chromaffin cells were obtained from the adrenal glands of adult (11-13 weeks old) WT, *Lyz1^−/−^* and *Nod1^−/−^* male mice. Amperometric recordings were performed after 1-3 days in culture at room temperature. The bath solution contained: 140 mM NaCl, 2.7 mM KCl, 5 mM CaCl_2_, 1 mM MgCl_2_, 10 mM HEPES/NaOH, and 5-10 mM glucose. Osmolarity was around 300 mOsm, and the pH was adjusted to 7.3 with NaOH. The working potential of the carbon fiber electrode (CFE) was held at +700 mV with a patch-clamp amplifier (EPC10, HEKA, Germany). With a micromanipulator (MPC200, Sutter Instruments, USA), the CFE was slowly advanced to the chromaffin cell of interest.

Data acquisition and analysis: The amperometric currents were recorded and analyzed using the Quantal Analysis program in Igor Pro (Version 6.2, WaveMetrics, Lake Oswego, USA), written by Drs. Mosharov and Sulzer (*Mosharov & Sulzer, 2005*). Data for off-line analysis were low-pass-filtered at 1 kHz. A level of 5× the RMS of noise at baseline was set as the threshold for signal detection.

### *Ex vivo* K^+^-stimulated catecholamine secretion assay

Cultured primary chromaffin cells were transfected following a published procedure (*Meunier et al., 2013*). Briefly, primary chromaffin cells were plated into imaging chambers and allowed to adhere for 3 hours. Complete medium was replaced with Opti-MEM medium, and the cells were transfected with 1 μg GRAB_NE_ using 2 μl Lipofectamine 2000. After 60 minutes, the mixture of Opti-MEM-Lipofectamine-plasmid was replaced again by complete medium to end the transfection procedure. The cells were used for secretion assay 48 hours later. In the experiment with chromaffin cells from *Lyz1^−/−^* mice, iE-DAP (1 μg/ml) was added into the complete medium 6 hours prior to the secretion assay.

Cultured chromaffin cells from WT, *Lyz1^−/−^, Nod1^−/−^ Dbh-cre;Nod1^f/f^* and *Nod1^f/f^* mice expressing the GRAB_NE_ sensor were imaged using an inverted Ti-E A1 confocal microscope. 525/50 nm and 600/30 nm emission filters were used to collect the GFP and RFP signals. Chromaffin cells expressing the GRAB_NE_ sensor were first bathed in Tyrode’s solution and imaged during the course of the K^+^ stimulation. The change in fluorescence intensity of the GRAB_NE_ sensor was calculated by recording the changes in the green and red fluorescence. For the recording, the exposure time was set at 50 ms per frame and the acquisition frequency was imaging every 5 seconds to reduce light bleaching. The solution used for perfusion was the physiological solution Tyrode’s solution (adjusted to pH=7.3-7.4), and KCl was added to reach a concentration of 90 mM. The average fluorescence intensity at the five time points before K^+^ stimulation was defined as F0. The instantaneous ΔF/F0 was calculated as (Ft - F0)/F0.

### Measurement of epinephrine in mouse plasma and tissue

Whole adrenal glands from WT, *Lyz1^−/−^, Nod1^−/−^, Dbh-cre;Nod1^f/f^, Nod1^f/f^* and ABX mice (ABX mice were prepared according to the published procedure (*Hill et al., 2010*)) were dissected on an ice bath. Subsequently, the isolated adrenal glands were weighed and homogenized with acetonitrile (30 μl for every 1 mg of tissue). Ascorbic acid (final concentration at 3 mg/ml) was added as antioxidant into the acetonitrile. After thorough homogenization, samples were subjected to sonication for 60 seconds to fully extract catecholamines. The adrenal gland homogenates were centrifuged at 12,000 rpm for 10 minutes at 4°C. Supernatants were collected and stored at −80°C before quantification.

Mice were subjected to restraint stress in a piping bag for 25 minutes or 1 hour at 8 am. A hole was made in the tip of the piping bag to enable air flow. Animals could move their heads but were unable to move their arms and legs. For the control group, the mice were placed in the home cage at the same time without food and water. Rectal temperature and blood glucose readings were taken before and after the restraint. Blood samples were collected into tubes containing 5 mM EDTA through orbital sinus bleeding. Subsequently, the samples were centrifuged and the plasma samples were stored at −80°C before analysis.

The concentration of epinephrine in tissue or plasma samples was measured using an Adrenaline Research Elisa kit (BA E-5100) from Labor Diagnostika Nord.

### Immunofluorescence staining of PC-12 cells or primary chromaffin cells

Cells grown on coverslips were washed with cold PBS and fixed in 4% PFA for 15 minutes at 4°C. Cells were then washed three times with PBS and blocked with antibody dilution buffer (PBS supplemented with 5% goat serum and 0.1% saponin) for 30 minutes. Cells were incubated with primary antibodies at 4°C overnight followed by fluorophore-conjugated secondary antibodies at RT for 2 hours. Samples were washed in PBS containing 0.1% Tween-20 three times for each step. Antibody dilution buffer was used throughout for antibody incubation steps. Coverslips were then counterstained and mounted onto slides in Fluoromount-G. Confocal images were obtained with a Zeiss LSM 710 imaging system under a 63x oil objective.

### Immunofluorescence (IF) staining of tissue sections

Murine tissues were fixed in 4% PFA/PBS overnight, dehydrated and embedded in paraffin. Tissue slices (6 μm in thickness) were mounted on positively charged glass and dewaxed. Antigen retrieval was performed by incubation in 0.01 M sodium citrate buffer (pH6.0) for 25 minutes in a boiling steamer. For immunofluorescence staining, slides were then blocked with normal goat serum blocking reagent for 30 minutes, followed by sequential incubation with primary antibodies at 4°C overnight and fluorophore-conjugated secondary antibodies at RT for 2 h. Slides were then counterstained with DAPI and mounted in Fluor mount-G. Confocal images were obtained with Zeiss LSM 710 or Nikon Ti NE imaging systems.

For immunohistochemistry staining, slides were then treated with an UltraSensitive TM S-P Kit instead. After sequential incubation with hydrogen peroxide (~ 15 minutes), blocking serum (~ 20 minutes), primary antibodies (overnight at 4°C), biotinylated secondary antibody (~ 20 minutes) and streptavidin-peroxidase (~ 20 minutes), immunoreactivity was visualized using diaminobenzidine (DAB). The slides were counterstained with hematoxylin, mounted, and observed with a light microscope (Nikon Eclipse 90i).

### Gene expression analysis using quantitative reverse-transcription PCR

RNA was extracted from dissected adrenal medulla and stored in Trizol at −80°C. RNA was then quantified and purified with a PrimerScript RT reagent kit. q-PCR reactions were performed with SYRB Premix Taq on a Rotor Gene 6000 thermal cycler in triplicate. The following thermal cycling conditions were used: 95°C for 30 seconds, followed by 40 cycles of 95°C for 5 seconds and 60°C for 34 seconds. The specificity of q-PCR was verified with melting curves for each PCR reaction. The level of target mRNA was determined by Delta-Delta Ct values between the target and loading control. Primers for q-PCR are listed in the Supplementary Information.

### Western blotting

The dissected adrenal medullas were homogenized in RIPA buffer (150 mM NaCl, 1% NP-40, 0.5% sodium deoxycholate, 0.1% SDS, 50 mM Tris pH8.0, 5.0 mM EDTA pH8.0, 0.5 mM dithiothreitol) supplemented with a protease inhibitor cocktail and 10 mM PMSF. The lysates were placed on ice for 30 minutes, then centrifuged at 12,000 rpm (10 minutes) to remove cell debris. Equal amounts of protein were separated on SDS-PAGE gels and blotted onto immobilon-p PVDF membranes. After blocking in 5% skim milk in PBS-T (0.5% Tween-20) for 1 hour at room temperature, membranes were incubated with primary antibody at 4°C overnight and secondary HRP-conjugated antibodies and visualized by using ECL detection reagent. Protein band intensity was quantified using ImageJ software.

### Electron microscopic analyses of DCGs in adrenal chromaffin cells

Isolated adrenal glands from euthanized mice were washed with cold PBS and cut into halves. Tissues were then fixed in 0.1 M sodium phosphate buffer (pH7.2) containing 2.5% glutaraldehyde and 2% paraformaldehyde at 4°C overnight. Dehydration was performed with an ethanol gradient followed by infiltration and embedding with a SPI-Pon 812 Epoxy Embedding Kit. Sections were cut on a Leica Ultracut Ultramicrotome (Leica EM UC6) and stained with uranyl acetate and lead citrate. The digital images were obtained under a transmission electron microscope (Hitichi H-7650) at 80 kV.

For morphometric analysis, the diameters and densities of typical and atypical dense core granules were measured using ImageJ software.

### Statistical analysis

Statistical analysis was performed using GraphPad Prism 8.1. Statistical tests were performed by GraphPad Prism. Student’s *t*-test was used for comparisons between two groups. One-way ANOVA was used for comparisons between more than two groups. The respective statistical test used for each figure is noted in the corresponding figure legends and significant statistical differences are noted as ns (*p*>0.05), \**p*≤0.05, \*\**p*≤0.01, \*\*\**p*≤0.001, \*\*\*\**p*≤0.0001.

## Acknowledgments

We thank Jiesi Feng and Yulong Li (State Key Laboratory of Membrane Biology, Peking University School of Life Sciences, Beijing) for pDisplay-NE1m-IRES-mCherry-CAAX plasmid, Cheng Zhan (National Institute of Biological Sciences, Beijing, China) for *Dbh Cre* mous*e* strain, Yihui Xu and the service station of CAS key laboratory of infection and immunity for technical support. This work was supported by National Key Research and Development Program of China (Grant no. 2017YFA0503403), the National Natural Science Foundation of China (Grant no. 31730028 and 81670482), the Instrument Development Project, CAS (YJKYYQ20180028 to J.S.), the National Natural Science Foundation of China (31527802 to J.S.) and Youth Innovation Promotion Association of the Chinese Academy of Sciences of China NO. 2017130 (P.C.).

## Supplementary Information

**Figure 1—figure supplement 1.**
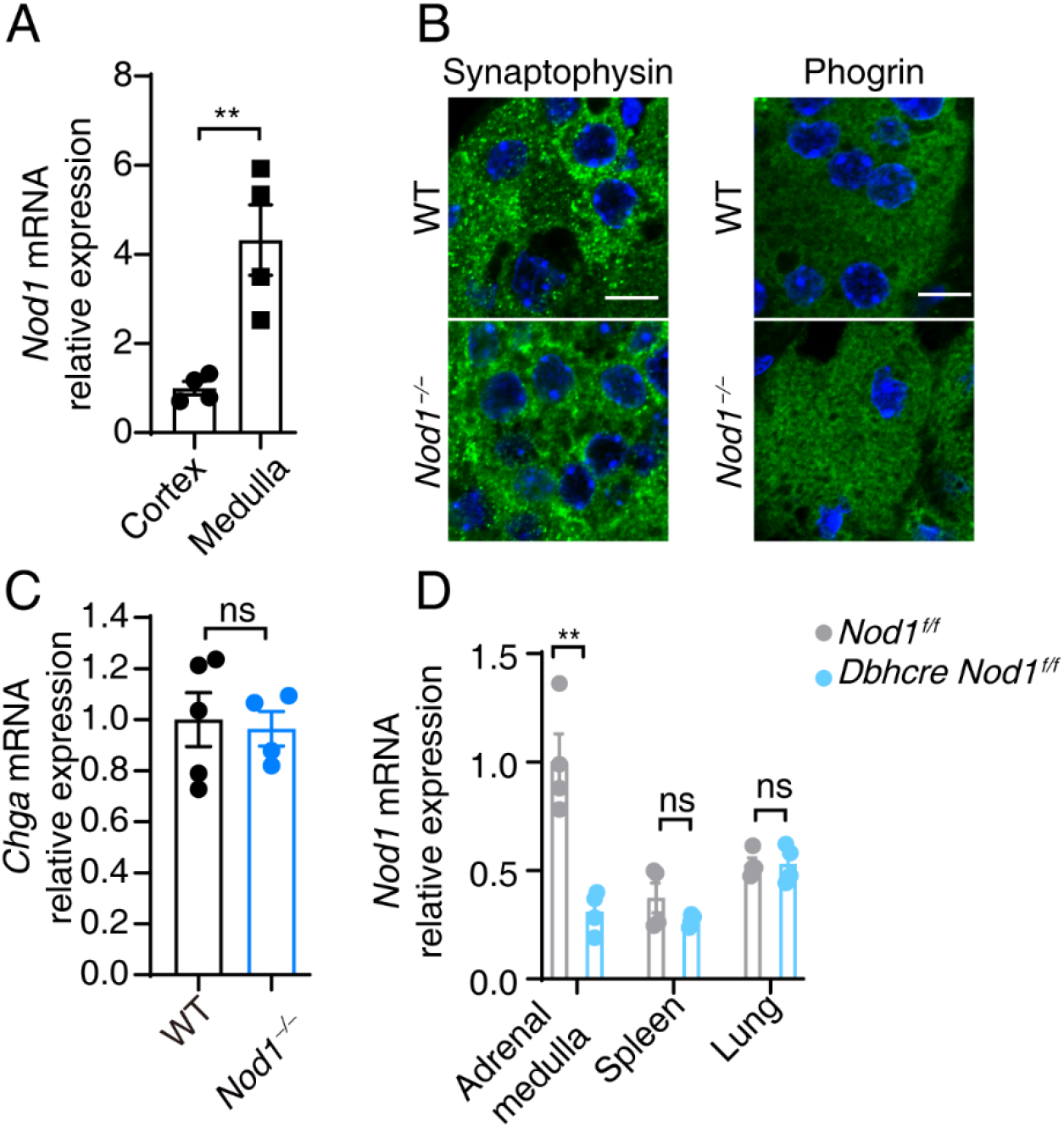
Nod1 is highly expressed in adrenal medulla. (**A**) Relative *Nod1^−/−^* mRNA in adrenal medulla and adrenal cortex determined by qPCR. The *GAPDH* mRNA level was used for normalization. (**B**) Immunostaining and confocal imaging of synaptophysin and phogrin in paraffin sections of adrenal gland from WT and *Nod1^−/−^* mice. (**C**) Relative *CHGA* mRNA level in adrenal medulla determined by qPCR. The *GAPDH* mRNA level was used for normalization. (**D**) Relative *CHGA* mRNA in adrenal medulla, spleen and lung of *Nod1*^f/f^ and *Dbh-cre;Nod1*^f/f^ mice determined by qPCR. The *GAPDH* mRNA level was used for normalization. Statistically significant differences were determined using Student’s *t*-test (**A, C**), and ANOVA (**B**). ns, indicates no significant difference (*p*>0.05). \*\**p*<0.01. Summary plots show mean with s.e.m. Each symbol represents the value from an individual animal.

**Figure 2—figure supplement 1.**
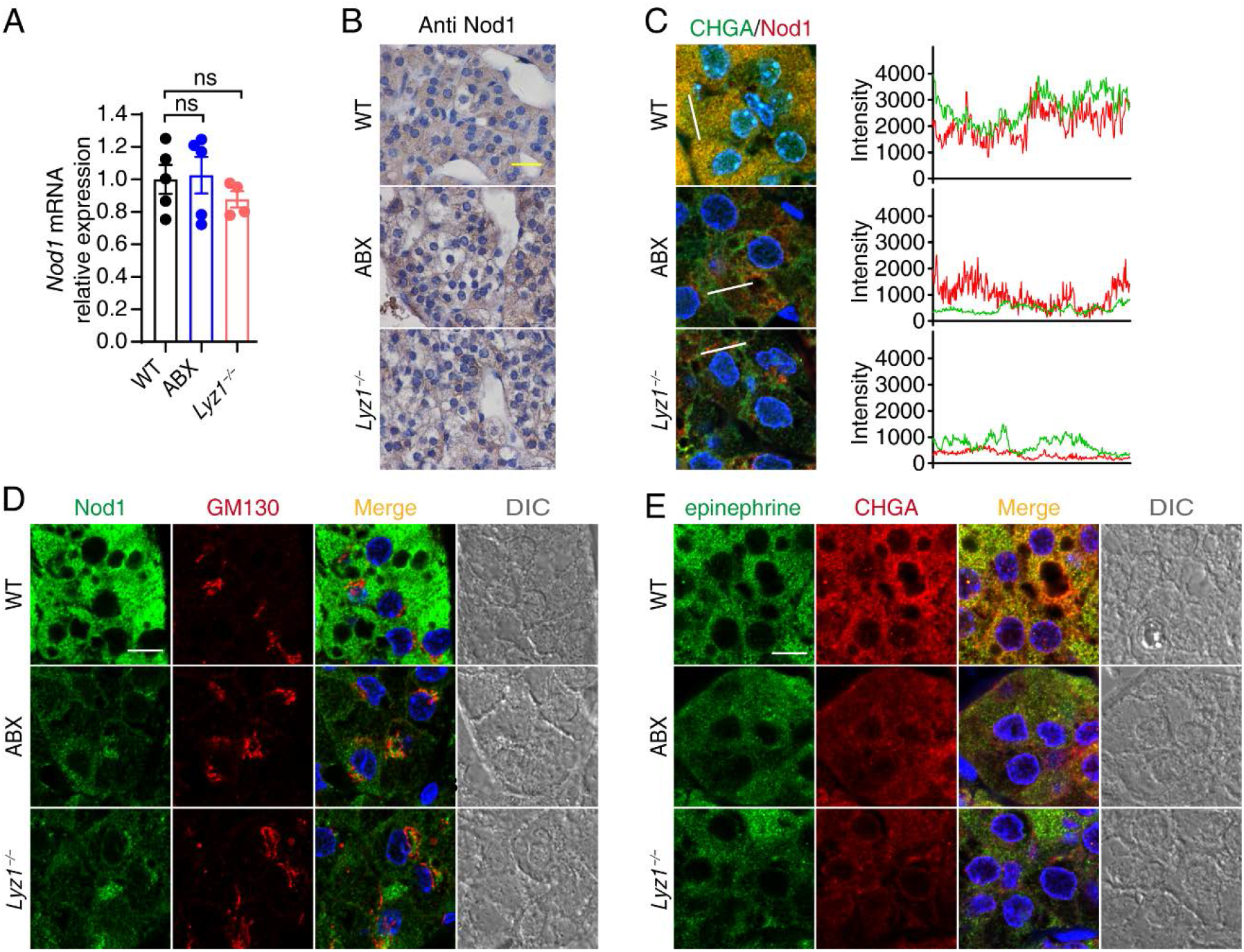
Nod1 localizes on DCGs to direct CHGA retention. (**A**) Relative *Nod1^−/−^* mRNA level in adrenal medulla from WT, ABX and *Lyz1^−/−^* mice. (**B**) Immunohistochemical staining (IHC) of Nod1 in paraffin sections of adrenal glands from WT, ABX and *Lyz1^−/−^* mice. (**C**) The RGB profiler plugin from ImageJ was used to plot colocalization of CHGA and Nod1 along the white lines drawn in the images, which correspond to the “Merge” images in Figure 2A. (**D**) Immunostaining and confocal imaging of Nod1 (green) and GM130 (red) in paraffin sections of adrenal gland from WT, ABX and *Lyz1^−/−^* mice. (**E**) Immunostaining and confocal imaging of epinephrine (green) and CHGA (red) in paraffin sections of adrenal gland from WT, ABX and *Lyz1^−/−^* mice. Statistically significant differences were determined using ANOVA (*A*). ns, indicates no significant difference (*p*>0.05). Each symbol represents an individual mouse. Mean and s.e.m. are indicated.

**Figure 5—figure supplement 1.**
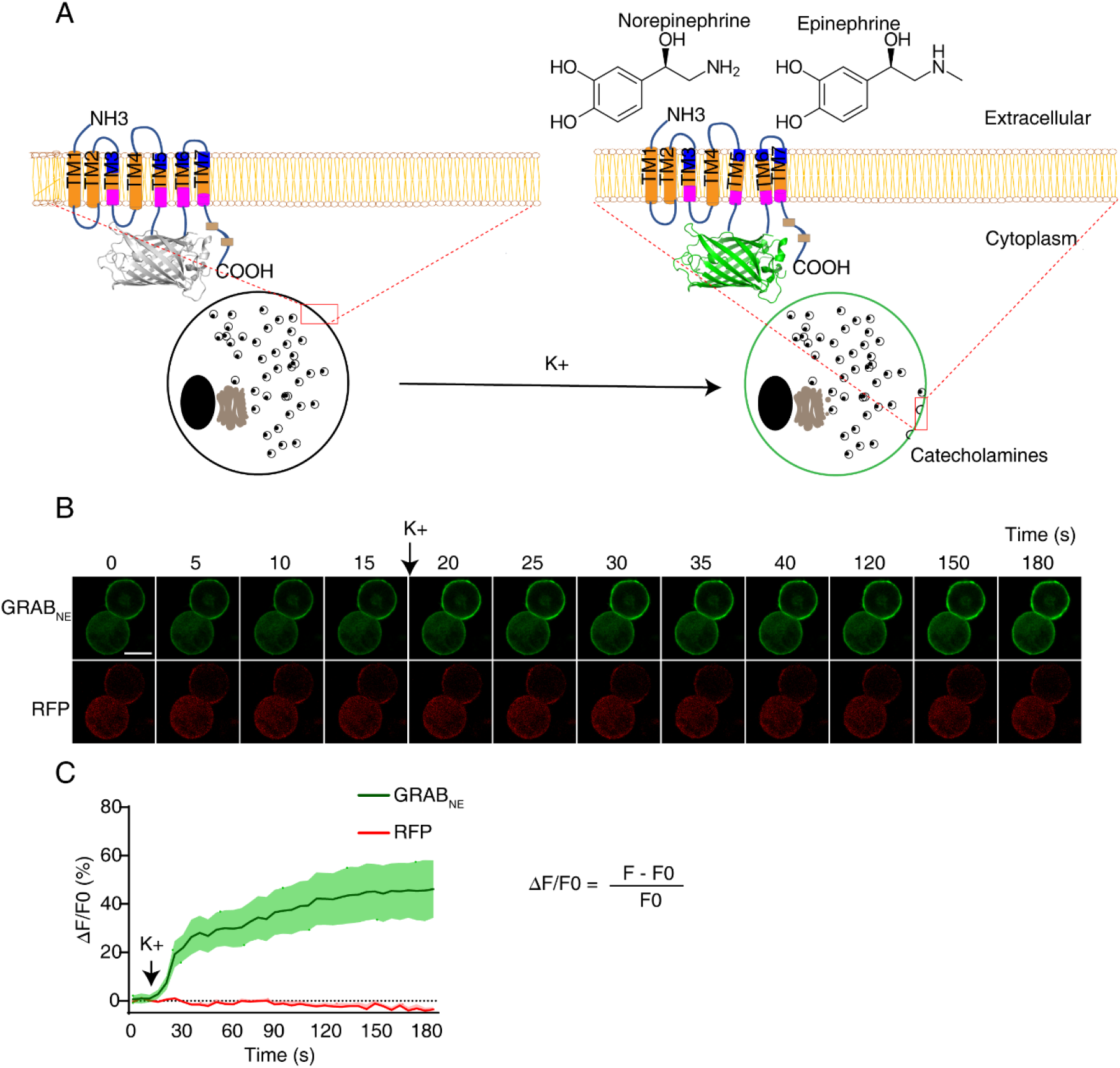
Epinephrine (norepinephrine) assay with GRAB_NE_. (**A**) Schematic diagram of the *ex vivo* K^+^-stimulated catecholamine secretion assay. (**B**) Representative images of green fluorescence of GRAB_NE_ and red fluorescence of RFP in transfected chromaffin cells during the course of treatment. (**C**) The recorded fluorescence changes (ΔF/F0) of GRAB_NE_ and RFP over the time course of the experiment (n= 5 cells from 3 mice). Scale bars,10 μm.

### Materials

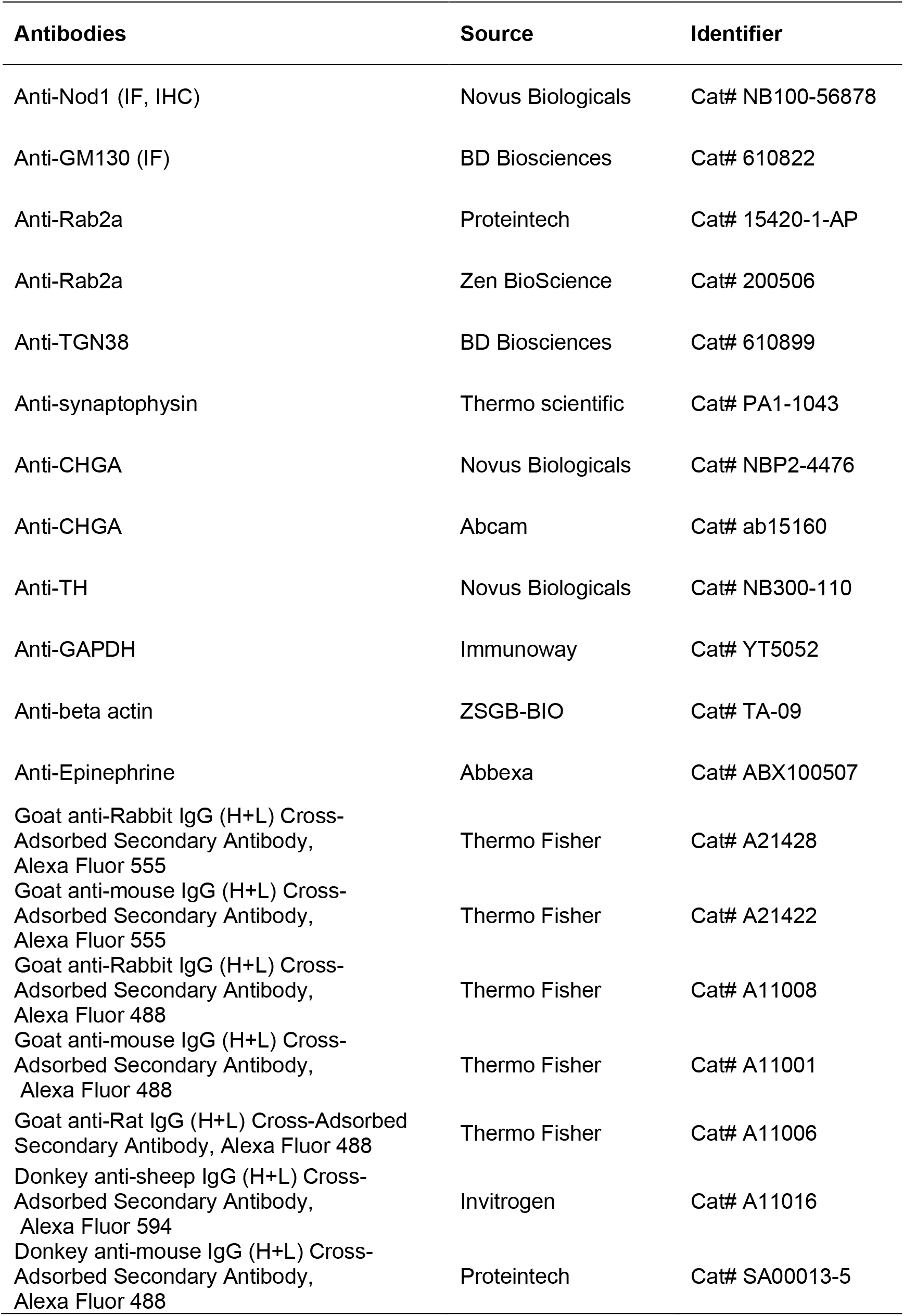
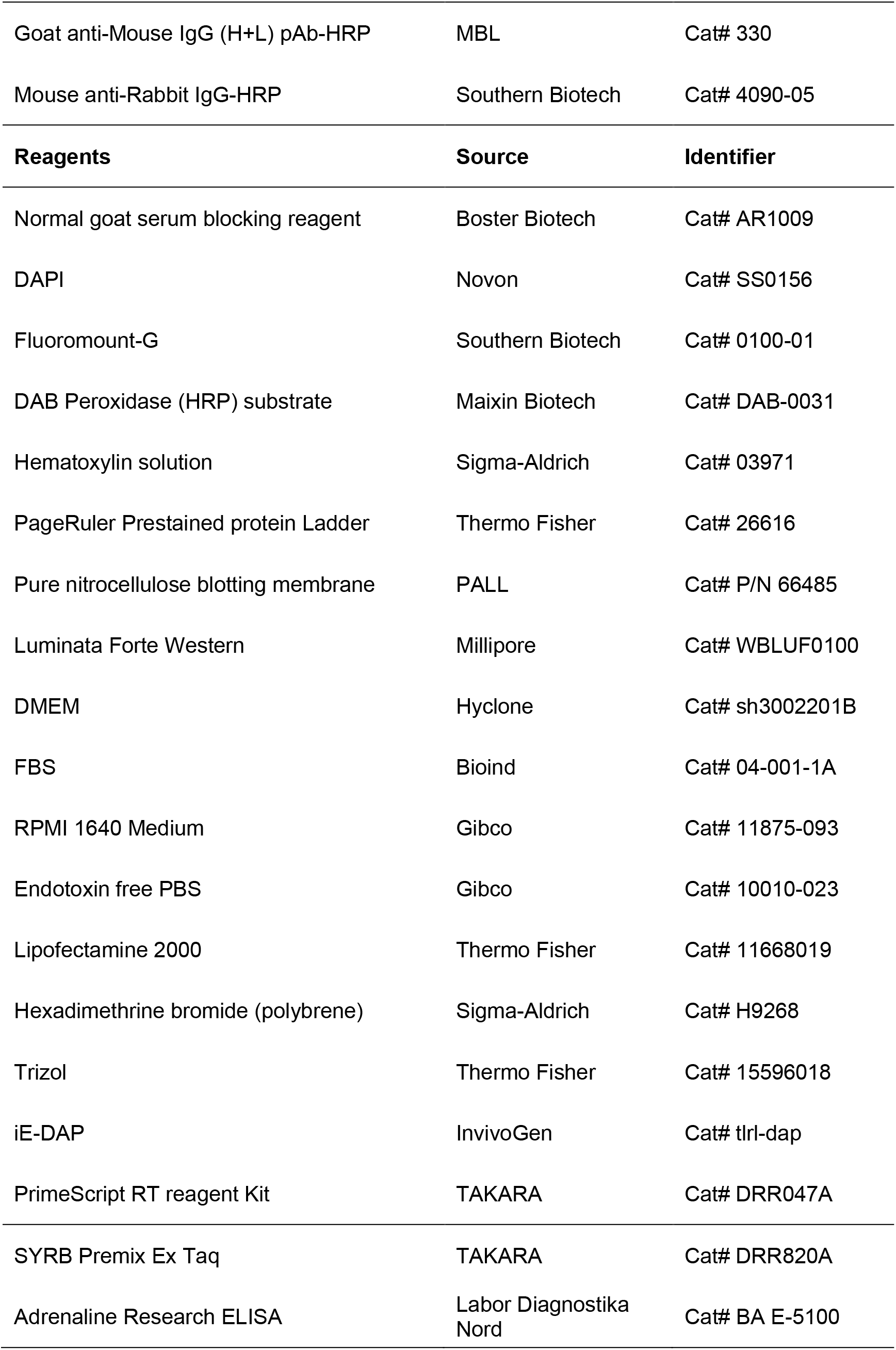
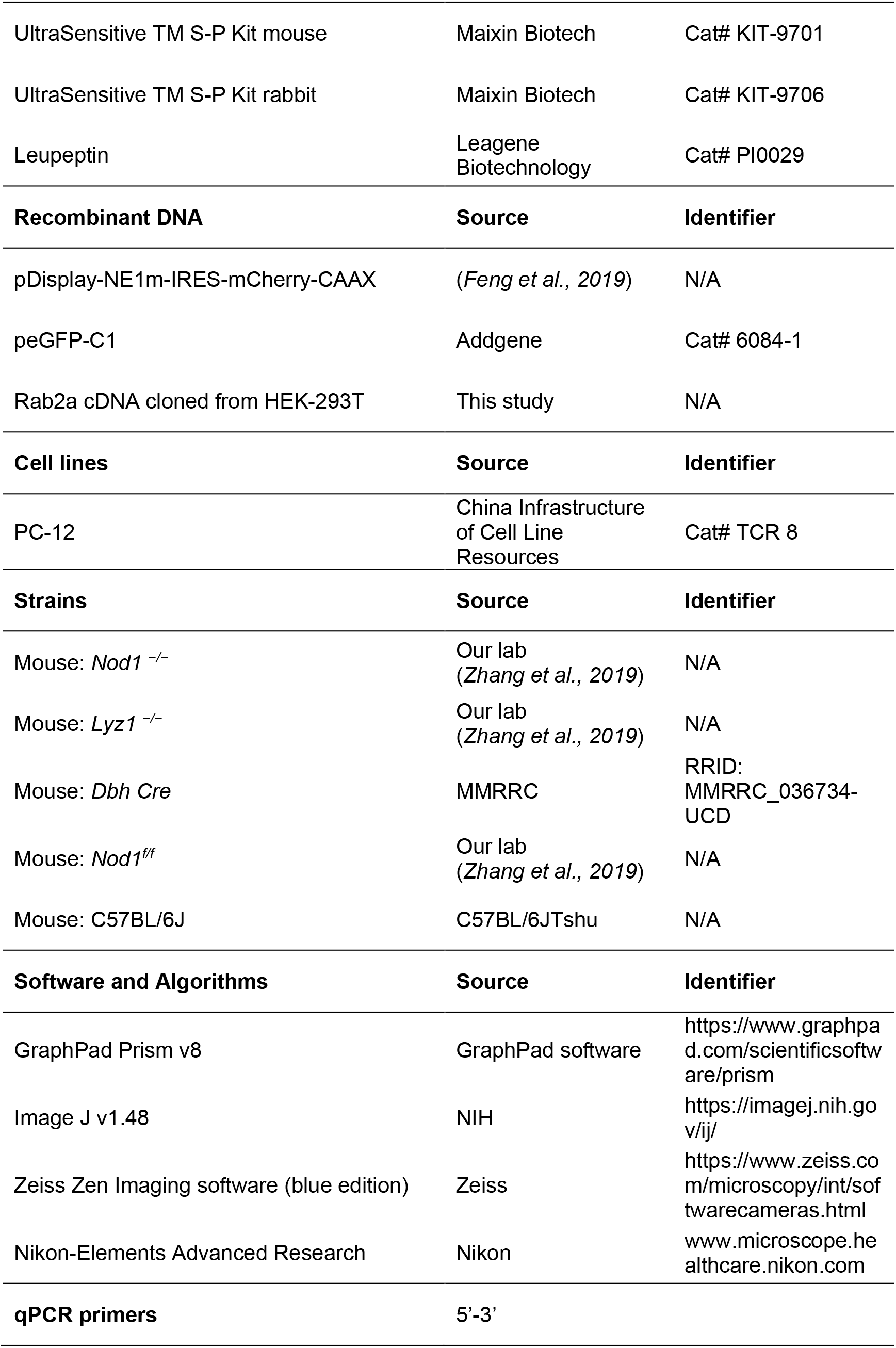
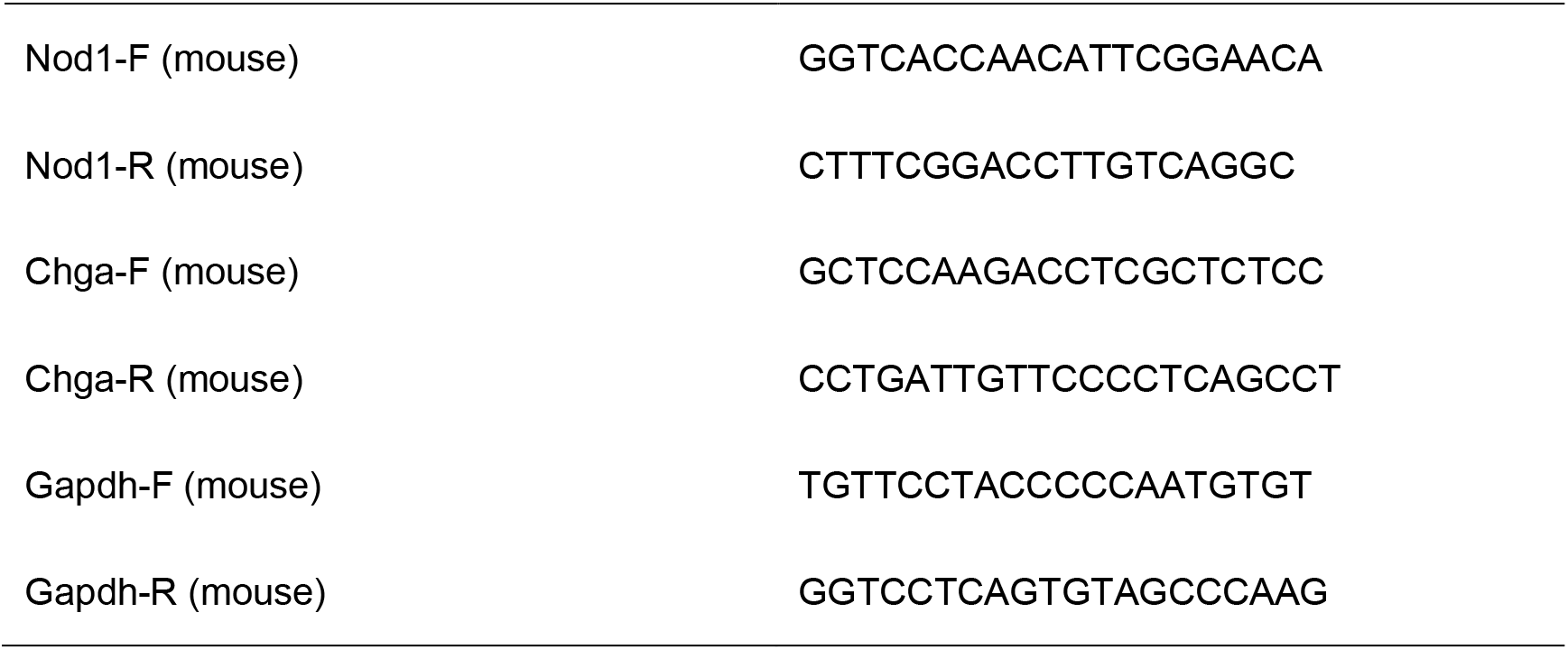

